# Structural comparative modeling of multi-domain ΔF508 CFTR

**DOI:** 10.1101/2021.11.17.468921

**Authors:** Eli Fritz McDonald, Hope Woods, Shannon T. Smith, Minsoo Kim, Clara T. Schoeder, Lars Plate, Jens Meiler

**Author notes:** **Corresponding author:** Jens Meiler.

## Abstract

Cystic Fibrosis (CF) is a common genetic disease caused by mutations in the Cystic Fibrosis Transmembrane Conductance Regulator (CFTR), an epithelial anion channel expressed in several vital organs. Absence of functional CFTR results in imbalanced osmotic equilibrium and subsequent mucus build up in the lungs - which increases the risk of infection and eventually causes death. CFTR is an ATP binding cassette (ABC) transporter composed of two transmembrane domains (TMDs), two nucleotide binding domains (NBDs), and an unstructured regulatory domain. The most prevalent patient mutation is the deletion of F508 (ΔF508), making ΔF508 CFTR the primary target for current FDA approved CF therapies. However, no experimental multi-domain ΔF508 CFTR structure has been determined and few studies have modeled ΔF508 using multi-domain WT CFTR structures. Here, we used cryo-EM density data and Rosetta comparative modeling (RosettaCM) to compare a ΔF508 model with published experimental data on CFTR NBD1 thermodynamics. We then apply this modeling method to generate multi-domain WT and ΔF508 CFTR structural models. These models demonstrate the destabilizing effects of ΔF508 on NBD1 and the NBD1/TMD interface in both the closed and open conformation of CFTR. Furthermore, we modeled ΔF508/R1070W and ΔF508 bound to the CFTR corrector VX-809. Our models reveal the stabilizing effects of R1070W and VX-809 on multi-domain models of ΔF508 CFTR and pave the way for rational design of additional drugs that target ΔF508 CFTR for treatment of CF.

**Author Summary:** Protein’s three-dimension shape determines their function, so when genetic mutation compromises the shape of vital proteins, it may cause disease. Such is the case in Cystic Fibrosis, a chronic genetic disease caused by mutations in the protein Cystic Fibrosis Transmembrane Conductance Regulator. Here, we work backwards from the shape of the wild-type protein – found in healthy people, to computationally model the shape of the most common Cystic Fibrosis mutant. Our computer models reveal distinct defects in the shape of the mutant Cystic Fibrosis Transmembrane Conductance Regulator protein in the area surrounding the mutation. We also model an important FDA approved Cystic Fibrosis drug, VX-809, into the mutant protein structure and show how VX-809 stabilizes the protein around the location of the mutation. The method we developed will pave the way for computational drug design for Cystic Fibrosis.

## INTRODUCTION

Cystic Fibrosis (CF) is caused by mutations in the cAMP-regulated, phosphorylation gated anion channel Cystic Fibrosis Transmembrane Conductance Regulator (CFTR) (1). CFTR is an ATP binding cassette type C (ABCC) transporter composed of two nucleotide binding domains (NBDs), two transmembrane domains (TMDs), and a flexible regulatory domain (2). CFTR undergoes a complex domain-domain assembly (**Figure 1A**) during biogenesis and folding. Deletion of phenylalanine 508 in NBD1 (ΔF508) is observed in 70% of patient alleles (3) and thus represents the most common cause of CF and target for drug development. ΔF508 destabilizes CFTR resulting in premature degradation and gating malfunction (4). Thus, the CF patient phenotype lacks CFTR mediated anion transport at the epithelial apical plasma membrane in the lung epithelia, hindering osmotic regulation and preventing cilia at the tissue/air interface from recycling mucus. Mucus build-up leads to poor lung function, is prone to infection, and ultimately leads to death (5).

**Figure 1.**
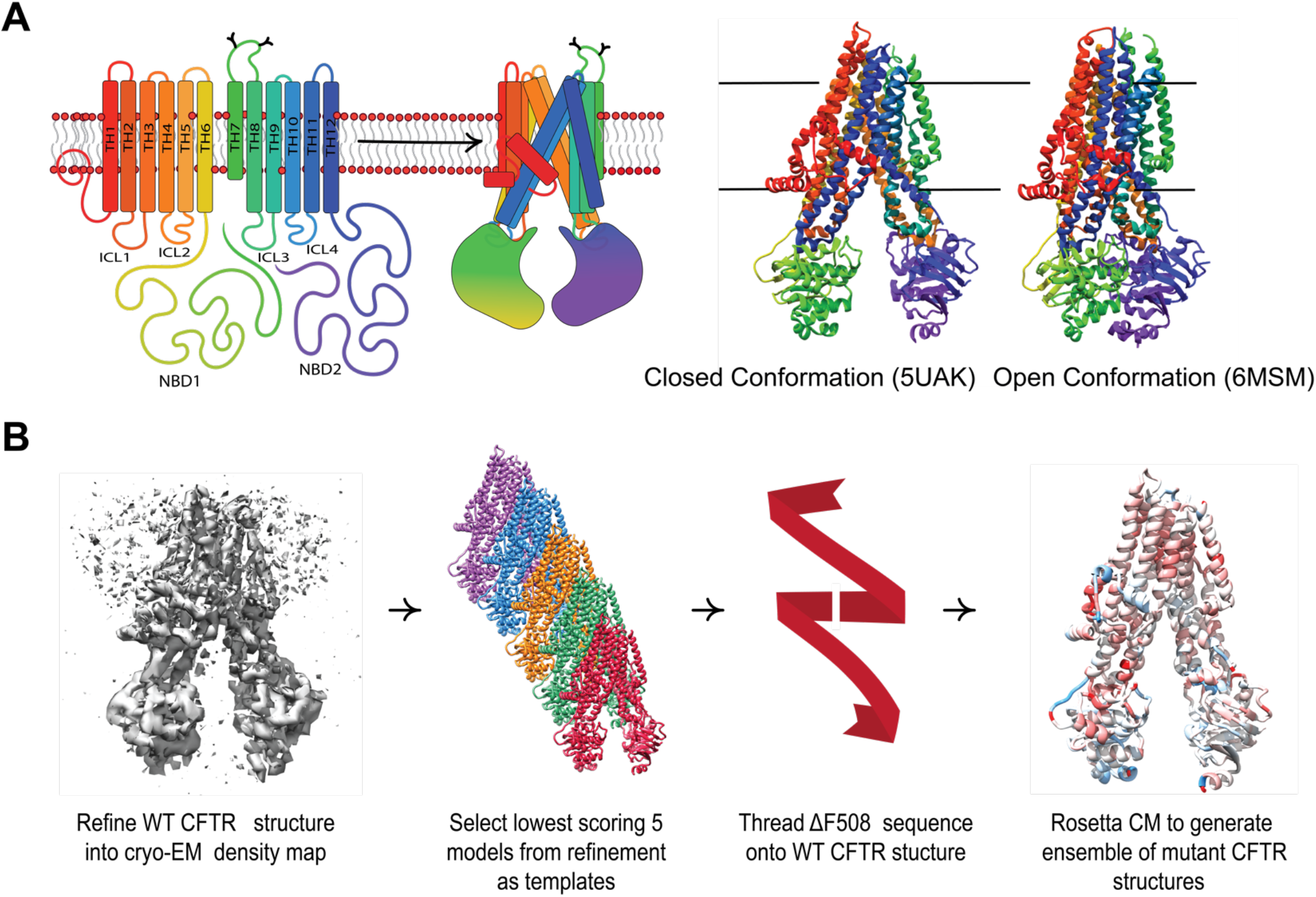
Comparative modeling captures multi-domain CFTR thermodynamics. **A**. The complex topology of CFTR involves interdomain contacts formed during the folding process that include intercellular loops (ICLs) interfacing with the cytosolic NDBs. The closed PBD ID 5UAK(2) (left) and open PBD ID 6MSM(20) (right). **B**. Our workflow for generating ensembles of ΔF508 models in this study (see Methods).

At present, CF treatment includes channel gating potentiation and CFTR folding correction through small molecules called potentiators and correctors, respectively (6,7). However, these compounds may interfere with birth control (8), cause testicular pain (9), and results in mental health side effects such as depression and psychotic symptoms (10). Understanding the atomic level mechanisms of CFTR correctors can facilitate computational design of improved CF therapeutics with fewer side effects. Cryo-EM (11) and computational modeling (12) revealed the binding site of FDA approved corrector VX-809 to WT CFTR. However, VX-809 binding to its primary target in the clinic, ΔF508 CFTR, remains poorly understood.

VX-809 stabilizes ΔF508 at the NBD1/TMD interface (13,14) and importantly, ΔF508 requires both NBD1 and NBD1/TMD interface correction to function (15,16). Thus, understanding the structural effects of ΔF508 on NBD1 and the NBD1/TMD interface with atomic resolution offers a basis for rational, structure-based drug design. Previous studies have used NBD1 crystal structures to simulate ΔF508 and understand the atomic effects on this single domain (17–19). However, despite the importance of the NBD1/TMD interface for ΔF508 correction, few studies leveraged recently published multi-domain CFTR structures to model ΔF508 CFTR (2,20– 22). Furthermore, no experimental structures of ΔF508 CFTR have been determined to date.

Here, we used Rosetta to model WT and ΔF508 CFTR. We first refined an ensemble of WT CFTR Rosetta models into the cryo-EM density (23) and then used the lowest scoring refined models as templates for Rosetta comparative modeling (RosettaCM) (24) to model ΔF508. We tested the optimal template number against thermodynamic data published on CFTR second site suppressor (SSS) NBD1 mutants (15,16,25). Next, multi-domain WT and ΔF508 CFTR structures including TMD1, NBD1, TMD2, and NBD2 were modeled using RosettaCM. We discussed our results in the context of abundant biochemical information about ΔF508 CFTR folding (26–28). Our models successfully captured ΔF508 CFTR thermodynamic destabilization consistent with folding defects in NBD1 and at the NBD1/TMD interface. Next, we modeled ΔF508/R1070W CFTR and demonstrated its ability to stabilize the NBD1/TMD interface. Finally, ΔF508 CFTR bound to VX-809 was modeled using RosettaCM and showed that drug binding decreased the total energy of the open state, but also stabilized the local region in the closed state. To our knowledge, this study presents the first attempt to model the multi-domain ΔF508 CFTR protein *in silico* using methods compatible with computer-aided drug design in the Rosetta Software Suite, a first step towards rational drug design for CF treatment.

## RESULTS AND DISCUSSION

### Refining CFTR models into available Cryo-EM Density Data

To model ΔF508, we sought to effectively sample CFTR conformational space *in silico* and generate a biophysically realistic set of template structures. This is needed as available CFTR structures are well determined in the TMDs (2.4 Å) but poorly determined in the NBDs (4.8-6Å) resulting in low average resolutions ranging from 3.2 to 3.9 Å (2,20–22). This motivated us to refine the WT structures into the cryo-EM density maps according to a previously established approach (23). Refinement generated a diverse set of models that sample the conformational space inherently accessible in the cryo-EM density map (23). For RosettaCM (29), we chose a subset of refined models to use as templates.

After optimizing refinement parameters for structural diversity (**Supplemental Figure S1**, see Methods), we refined 2000 models into the cryo-EM density maps (23) for the human dephosphorylated/closed conformation (PDB ID 5UAK) (2) and the human phosphorylated/open conformation (PDB ID 6MSM) (20). To evaluate the WT CFTR ensemble diversity, we calculated the Cα per-residue root mean squared deviation (RMSD) for each conformation from the respective published structure. Next, we mapped the average Cα per-residue RMSD onto the respective CFTR model from 0-4 Å to demonstrate visually which regions of CFTR show higher RMSD and are thus interpreted as inherently more flexible (**Figure 2A, 2B**). The ensemble demonstrated no substantial change in flexibility after 1000 models had been generated. Thus, we chose to stop generating models after model 2000 assuming a good sampling of the available conformational space.

**Figure 2.**
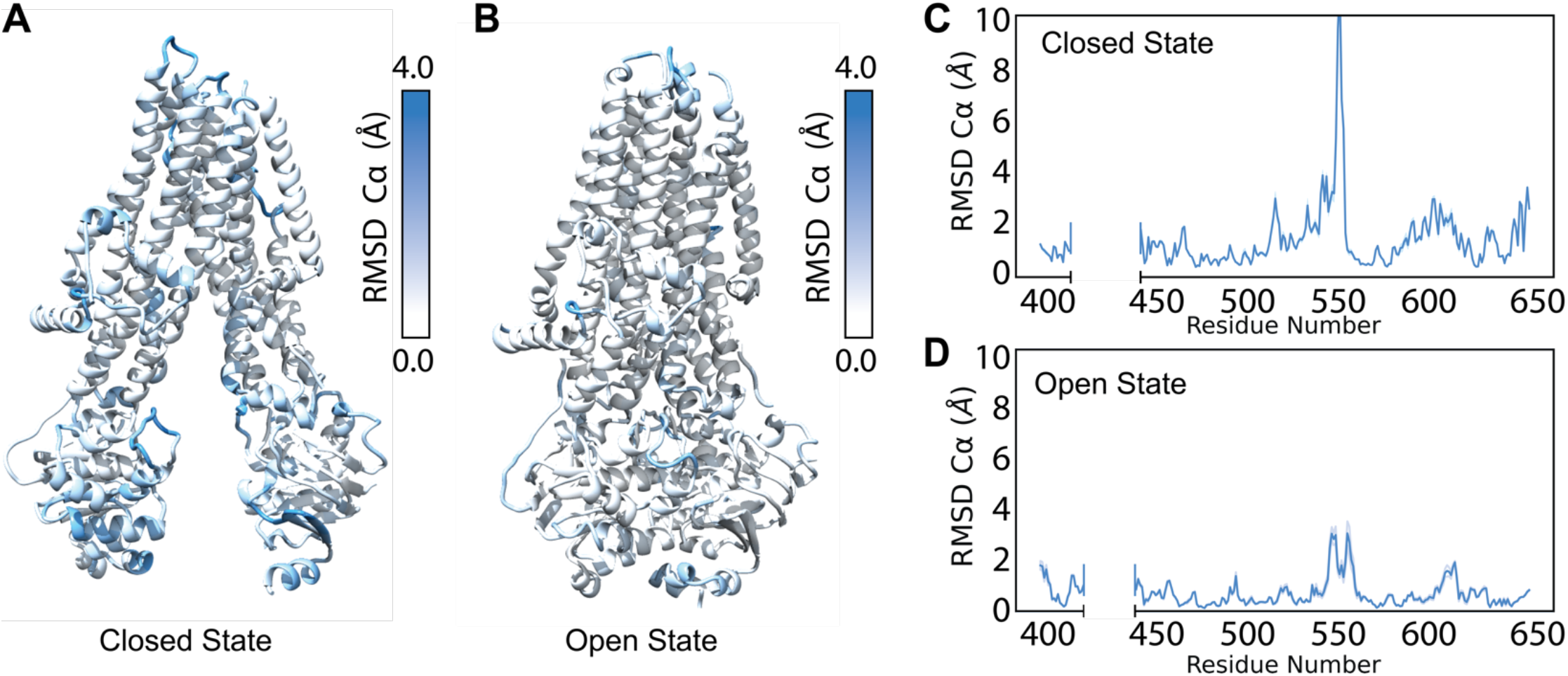
Refinement into the cryo-EM density generates a diverse ensemble of structures. **A**. Average RMSD of the best scoring (lowest 10% by potential energy function) 5UAK cryo-EM refinement models mapped onto 5UAK. **B**. Average RMSD of the best scoring (lowest 10% by potential energy function 6MSM cryo-EM refinement models mapped onto 6MSM. **C**. The average NBD1 RMSD of the best scoring 100 5UAK refinement models. The blue shading represents a 95% confidence interval, and the large RMSD demonstrates high structural diversity in the SDR (residues 526-547). **D**. The average NBD1 RMSD of the best scoring 100 6MSM refinement models. The blue shading represents a 95% confidence interval.

Overall, the poorly determined NBDs showed greater structural diversity than the TMDs, as measured by RMSD from the published model (**Figure 2A, 2B**). Likewise, in the closed state, NBDs showed greater structural diversity than the open state (**Figure 2A, 2B**). This likely resulted from the dimerized NBDs in the open state, which increased stability and lead to higher resolution cryo-EM data. Thus, the closed conformation offered a greater sampling of conformational space in the refinement ensemble than the open state. The refinement resulted in an ensemble of CFTR models with diverse conformations of loop regions such as extracellular loops and NBD1 loops.

We plotted the Cα per-residue RMSD for NBD1 in the open and closed state to compare which sub-domains and regions demonstrated greater structural diversity between the conformations (**Figure 2C, 2D, Supplemental Figure S2A, S2B**). Notably, the structurally diverse region (SDR), residues 527-547, showed substantial increased Cα RMSD in both conformations, consistent with the known flexibility of the region (**Figure 2C, 2D**) (30). The loop connecting H7 and β-strand 9 (S9) (residues 595-605) also showed flexibility, consistent with previous NBD1 only MD simulations indicating that S9 unfolds early in NBD1 unfolding (**Figure 2C, 2D**) (31). Together, these data suggest our cryo-EM refinement ensemble successfully captured the conformational flexibility of CFTR consistent with previous computational and experimental studies.

### Testing ΔF508 Modeling with CFTR NBD1 Second Site Suppressor Mutations

We sought to accurately model ΔF508 CFTR by leveraging the models generated during cryo-EM refinement. The low resolution cryo-EM data provided information on the conformational space inherently accessible to WT CFTR, although it remains unclear if ΔF508 CFTR samples the same conformational space. Given a novel sequence, RosettaCM samples the conformational space of homologous models called *templates* (24). Instead of a novel sequence and homologous models, we used the ΔF508 sequence and WT cryo-EM refinement models with the lowest potential energy scores as templates (see Methods).

We restricted our simulations to NBD1 (residues 385-402 and 435-644) because experimental CFTR thermodynamic data are only available for NBD1 (15,16,25) (**Supplemental Table 1**). Considering all residues with determined coordinates from the closed (5UAK) and open (6MSM) state NBD1 (residues 385-402 and 439-637), these regions superimposed well with an RMSD of 2.23 Å, lower than the published resolution of either structure (2,20) (**Supplemental Figure S3A**). Thus, we chose to test only the closed state NBD1 as the lower resolution offers more conformational sampling and the two structures are similar.

We generated ΔF508 models by threading the ΔF508 fasta sequence onto the WT model. The gap can be closed without major perturbation of the structure of NBD1 (**Figure 3A**). Deletion of F508 prematurely terminated helix 3 (H3) causing the loop connecting H3 and H4 to shift. This is consistent with the loop shift observed experimentally in the ΔF508 NBD1 crystal structure (32). Likewise, I506 and I507 side chains remained in their location when compared to WT (**Figure 3A)**. Furthermore, G509 was pulled closer to H3 but fails to form a backbone hydrogen bond with I507. The V510 side chain moved slightly, tightening the loop similar to the ΔF508 NBD1 crystal structure (32). We included ATP at the degenerate binding site because NBD1 is known to fold with ATP as a scaffold at this site (33) (**Figure 3B**). Further, we simulated WT and ΔF508 CFTR NBD1 with stabilizing mutations called second site suppressor (SSS) mutations. We included NBD1 SSS mutations F494N, F494N/Q637R, V510D, I539T, and G550E/R553Q/R555K (**Figure 3B**) because experimental CFTR thermodynamic data are available for these SSS (15,16,25).

**Figure 3.**
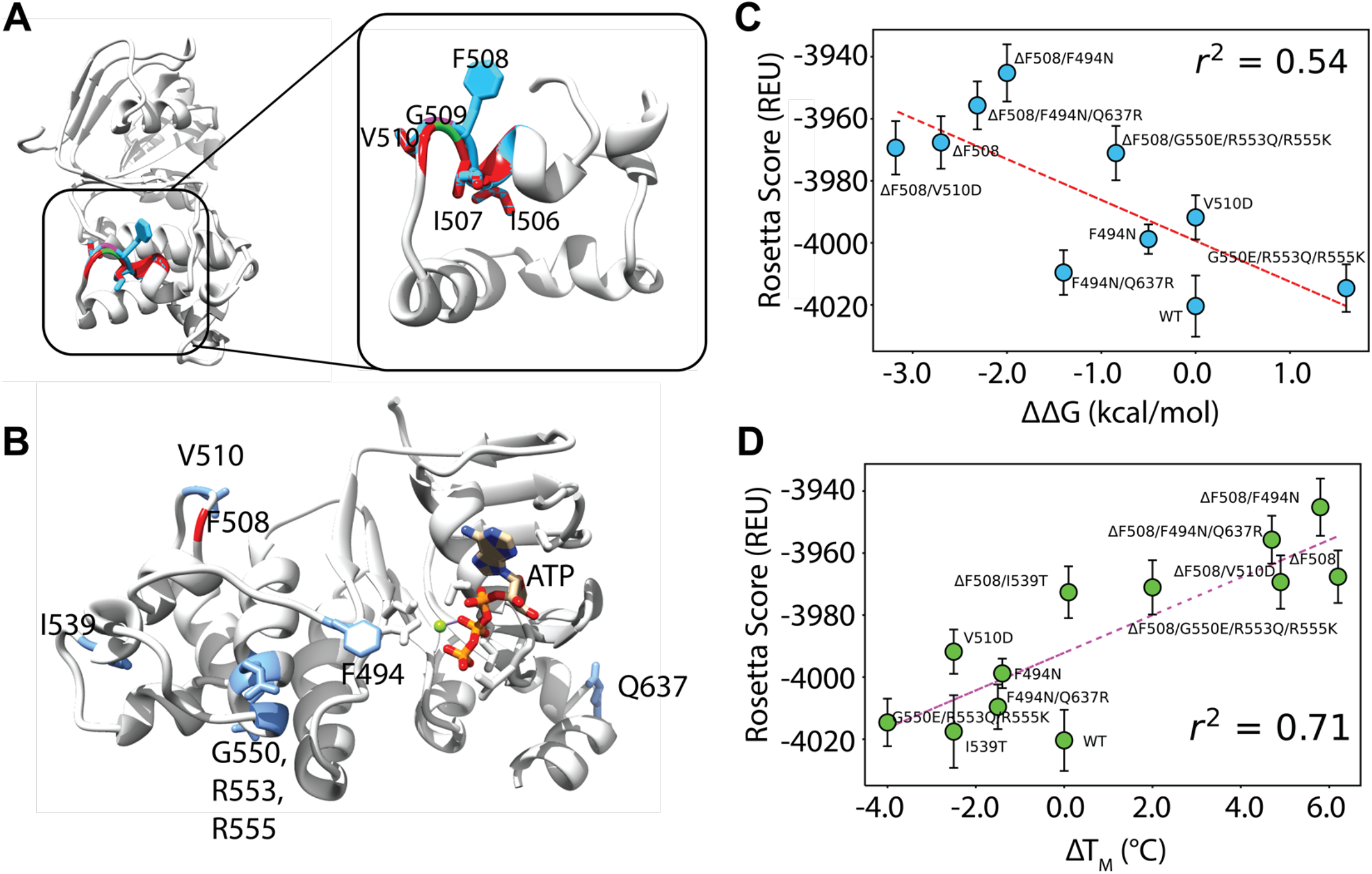
Comparative modeling of ΔF508 NBD1 using five templates correlates well with experimental data. **A**. An overlay of WT and ΔF508 CFTR NBD1 structures at the H3/H4 loop. WT is depicted in blue and ΔF508 is depicted in red with just the α-helical subdomain shown for clarity. Deletion of F508 leaves surrounding residues I506, I507 and V510 relatively unaltered. **B**. The model for testing included NBD1 bound to ATP. Residues mutated in second site suppressor mutations are shown in blue including F494N, V510, I539, G550, R553, R555, and Q637. **C**. Testing correlation between Rosetta score given in REU and ΔΔG values from the literature (**Supplemental Table S1**). Error bars represent standard error of the mean. Error in experimental data likely ranges with +/- 1-2 kcal/mol. R squared represents Pearson correlation coefficient. **D**. Testing correlation between Rosetta score given in REU and ΔT_M_ values from the literature (**Supplemental Table S2**). Error bars represent standard error of the mean. Error in experimental data likely ranges with +/- 1-2 C. R squared represents Pearson correlation coefficient.

We used NBD1 experimental T_m_ and ΔG data to test the number of cryo-EM templates needed as input for RosettaCM to successfully capture differences in ΔF508 thermodynamic instability with and without SSS mutations. Specifically, ΔT_m_ and ΔΔG with respect to WT NBD1 in each study were used to account for distinct experimental conditions between studies (**Supplemental Table 1**). We generated 1000 models for each mutation and took the average Rosetta score of the lowest scoring 50 models – or lowest scoring 5% of models. Next, the Rosetta score versus the ΔT_m_ and ΔΔG were plotted for each SSS mutation and we calculated the Pearson correlation coefficient between the Rosetta Scores and the experimental values. Of note, the experimental ΔTm values correlated with the experimental ΔΔG values with an r^2^ of 0.78 which we subsequently assumed represents a good correlation, given the limitations of the experiment data (**Supplemental Figure S3B**).

We determined that using 3, 4, 5, 7, and 9 templates resulted in a Rosetta score-ΔT_M_ Pearson correlation coefficient of 0.22, 0.45, 0.71, 0.59, and 0.22 respectively and a Rosetta score-ΔΔG correlation of 0.14, 0.27, 0.54, 0,57, and 0.25 respectively (**Figure 3C, 3D, and Supplemental Figure S3C-G**). Thus, five templates offered the best correlation (**Figure 3C, 3D**).

### ΔF508 Destabilizes Closed and Open State of human CFTR

CFTR is unique among ABC transporters to function as a phosphorylation gated anion channel. ΔF508 CFTR gates inefficiently and requires potentiators such as VX-770 to stabilize the open conformation (34). Given the clinical importance of CFTR channel gating, we modeled ΔF508 in both the closed and open conformations.

We generated 2000 structure ensembles of WT and ΔF508 CFTR using RosettaCM for both the closed (PDB ID 5UAK) (2) and open (PDB ID 6MSM) (20) CFTR conformations. We examined the lowest scoring 100 models in terms of Rosetta score (Rosetta Energy Units or REU), which represented the best scoring 5% of the models generated. We plotted structural Cα RMSD (relative to the lowest scoring WT model) vs. Rosetta score to determine global structural changes among the mutant models (**Figure 4A, 4B**). WT and ΔF508 models showed distinct structural shifts as measured by RMSD (**Figure 4A**). Furthermore, the ΔF508 models showed higher energy in terms of REU (**Figure 4B**). These data suggest our models captured ΔF508 thermodynamic instability in both conformations.

**Figure 4.**
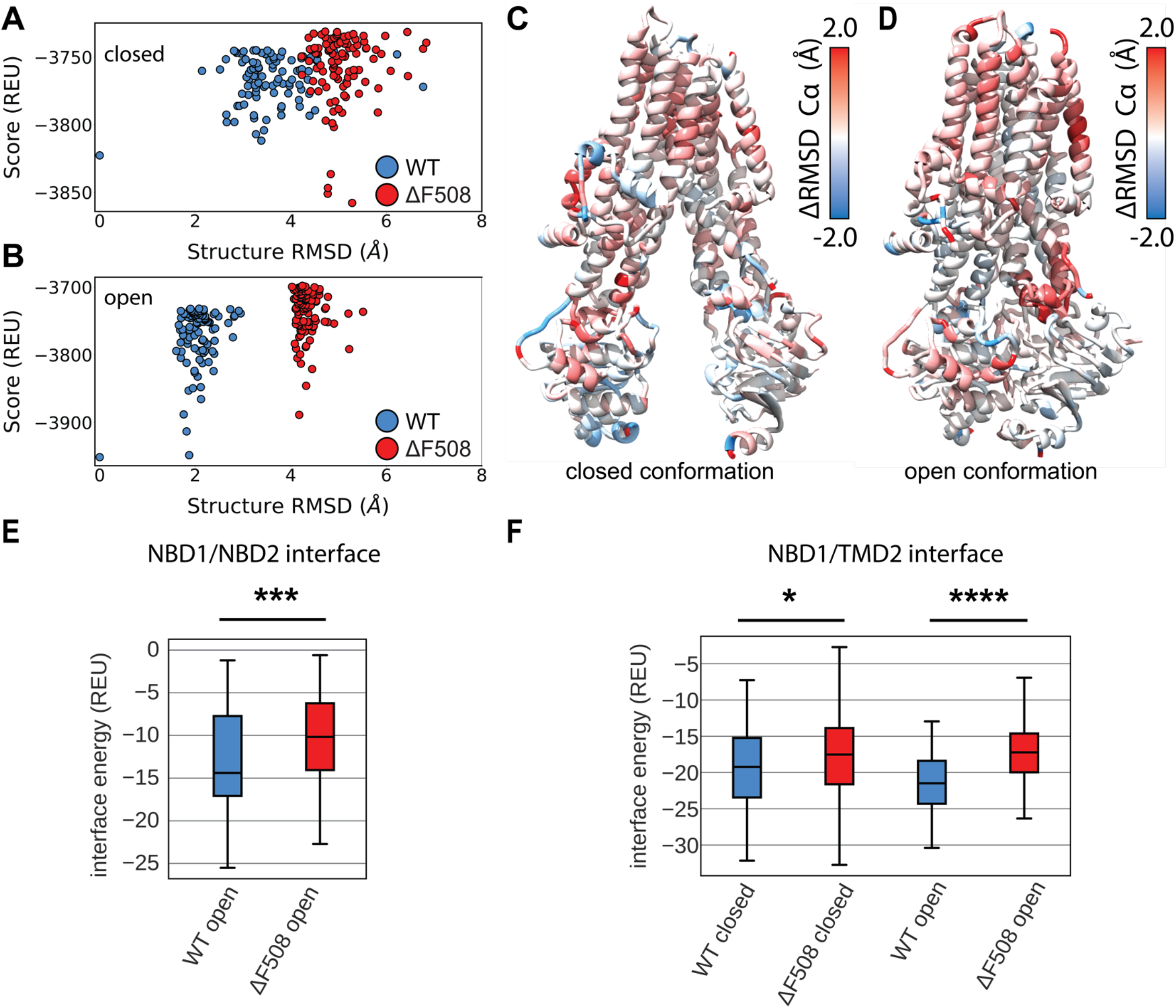
Comparative modeling of multi-domain ΔF508 CFTR shows thermodynamic instability and lose of interaction energy at key domain-domain interfaces. **A**. Cα RMSD vs. score plot of the lowest scoring 100 closed conformation models from ensembles of WT (blue) and ΔF508 (red) CFTR. RMSD is calculated relative to the lowest scoring WT model. Score is shown in REU. **B**. Cα RMSD vs. score plot of the lowest scoring 100 open conformation models from ensembles of WT (blue) and ΔF508 (red) CFTR. **C**. Average residue Cα RMSD of the lowest scoring 100 closed state WT models subtracted from the Cα RMSD of the lowest scoring 100 closed state ΔF508 models mapped on 5UAK. Here red represents region where the RMSD was higher in ΔF508 than WT, and blue represents regions where the RMSD was lower. **D**. Average residue Cα RMSD of the lowest scoring 100 closed state WT models subtracted from the Cα RMSD of the lowest scoring 100 closed state ΔF508 models mapped on 6MSM. **E**. Quantification of the residue-residue interactions at the NBD1/NBD2 interface across the lowest scoring 100 models. Only the open state is considered as the closed state lack the NBD dimer and hence there are no residue interactions to measure. **F**. Quantification of the residue-residue interactions at the NBD1/TMD2 interface across the lowest scoring 100 models.

We sought to determine where ΔF508 confers thermodynamic instability to the CFTR structure. We used the residue RMSD from the lowest scoring model in each ensemble as a surrogate for structural flexibility associated with thermodynamic instability. Hence, we compared the flexibility of the lowest scoring 100 models in each WT and ΔF508 ensembles in both conformations (**Supplemental Figure S4**). We subtracted the WT residue RMSD from the ΔF508 residue RMSD and mapped the difference onto the published closed and open conformations (**Figure 4C, 4D**). Here red represents regions where ΔF508 showed more flexibility than WT, and blue represents regions where ΔF508 showed less flexibility. By this metric, ΔF508 demonstrated higher flexibility for both conformations in the α-helical subdomain (residues 500-540), specifically in helix 4B following F508 (**Figure 4C, 4D and Supplemental Figure S4**). The intercellular loops (ICLs) also demonstrated higher RMSD in ΔF508, particularly ICL4 in the closed state (**Figure 4C, Supplemental Figure S4A**) and ICL2 in the open state (**Figure 4D, Supplemental Figure S4B**). These data suggest our multi-domain ΔF508 reproduces the known destabilizing effects particularly in NBD1 and the ICLs.

Finally, multi-domain CFTR models allowed us to examine the energetic changes at the domain-domain interfaces. We calculated the residue interaction energy between all residues in the structures. We plotted the interaction energies between domains for the best scoring 100 models in terms of REU (**Supplemental Figure S5**). Next, WT and ΔF508 were compared by summing the interaction energy across the interface and plotting the distribution of sums as boxplots (**Supplemental Figure S6A**). ΔF508 significantly increases the residue interaction energy between NBD1 and NBD2 in the open state (**Figure 4E**), consistent with the notion that ΔF508 drives NBD2 unfolding *in vivo* (27). Furthermore, ΔF508 significantly increases the interaction energy between NBD1 and TMD2 (**Figure 4F**), which has long been suggested to be the predominant folding defect of ΔF508 (15,16). Finally, ΔF508 significantly increases the interface energy between TMD1/NBD2 and TMD1/TMD2 in the open state as well (**Supplemental Figure S6B**). These data suggest our models captured the thermodynamic instability of the NBD1/NBD2 dimer interaction and the NBD1/TMD2 interface expected for the mutant consistent with destabilizing effects on these interfaces (15,16).

### Modeling ΔF508/R1070W in multi-domain CFTR lowers interactions energy at the NBD1/TMD2 interface

Deletion of F508 leaves the aromatic pocket in ICL4 formed by F1068, Y1073, and F1074 empty, but the CFTR SSS mutation R1070W introduces a tryptophan into this pocket rescuing folding (**Figure 5A**) (15,16). Interestingly, ΔF508/R1070W resisted correction by VX-809 indicating the SSS mutant and drug function via a similar mechanism stabilizing the NBD1/TMD interface (13). Thus, R1070W represents a clinically relevant SSS to study in the context of multi-domain CFTR structure. To further evaluate our multi-domain CFTR modeling approach, we simulated ΔF508/R1070W and examined its effect relative to WT and ΔF508 CFTR.

**Figure 5.**
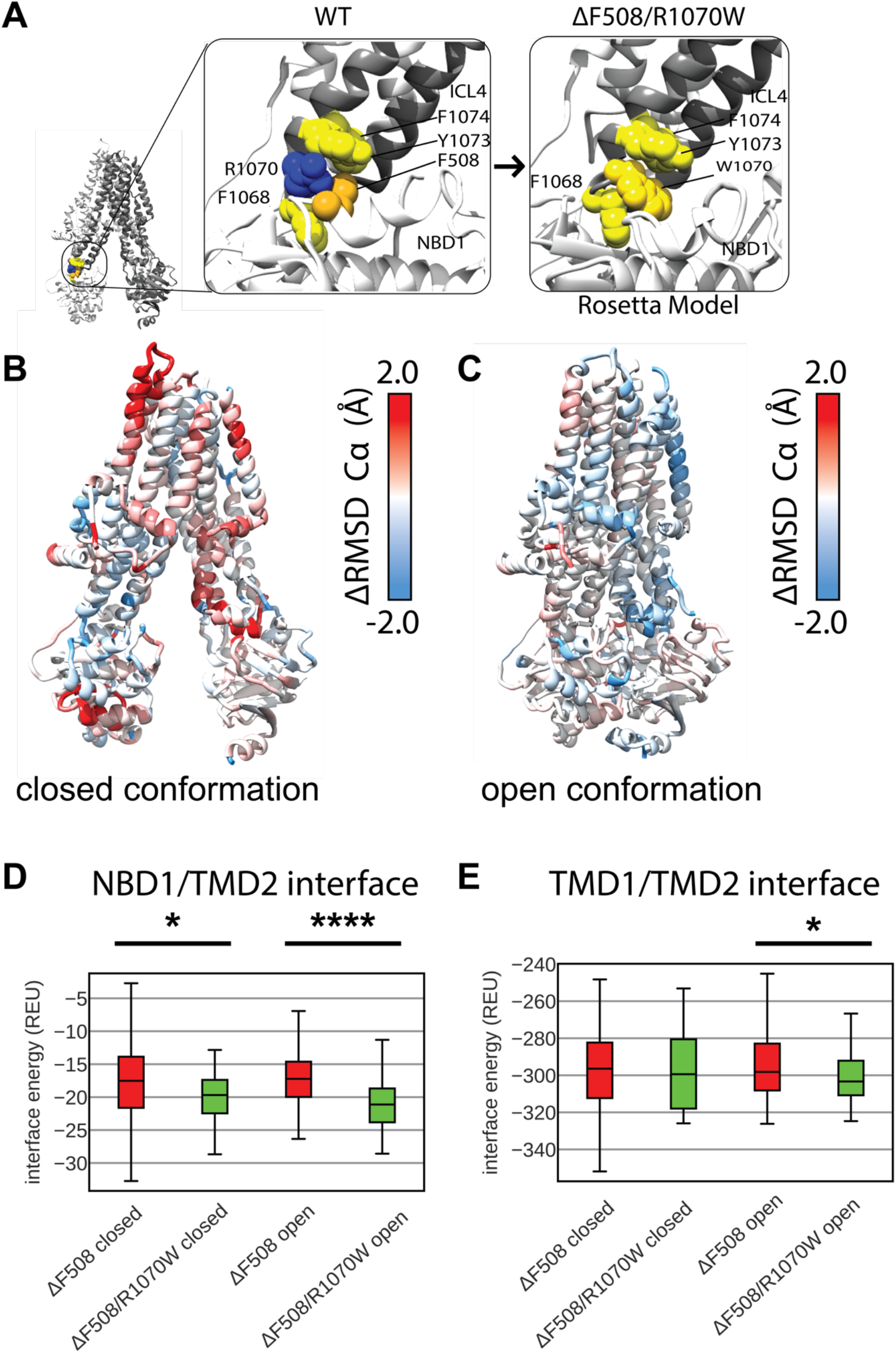
R1070W stabilizes the NBD1/TMD2 interface. **A**. F508 in 5UAK CFTR (gold spheres) contacts an aromatic pocket in ICL4 formed by F1068, Y1073, and F1074 (yellow spheres). This aromatic pocket is filled with R1070 (blue) is mutated to a tryptophan (left). **B**. Average residue Cα RMSD of the lowest scoring 100 closed state ΔF508 /R1070W models subtracted from the Cα RMSD of the lowest scoring 100 closed state ΔF508 models mapped on 5UAK. Here red represents region where the RMSD was higher in ΔF508/R1070W than ΔF508 alone, and blue represents regions where the RMSD was lower and hence stabilized by R1070W. **C**. Average residue Cα RMSD of the lowest scoring 100 closed state ΔF508 /R1070W models subtracted from the Cα RMSD of the lowest scoring 100 closed state ΔF508 models mapped on 6MSM. **D**. Quantification of the residue-residue interactions at the NBD1/TMD2 interface across the lowest scoring 100 models. **E**. Quantification of the residue-residue interactions at the TMD1/TMD2 interface across the lowest scoring 100 models. R1070W likely stabilize TMD2 enough to reduce the interaction energy between the TMDs in the open conformation.

We generated 2000 structure ensembles of ΔF508/R1070W using RosettaCM in both the closed and open CFTR conformations. We examined the best scoring 100 models (best scoring 5%) in terms of Rosetta Score. We determined the ensemble structural shift and thermodynamic changes conferred by R1070W by plotting structural RMSD vs. Rosetta score. ΔF508/R1070W shifts the ensemble structure very little from ΔF508 models as measured by RMSD and increases the energy (**Supplemental Figure S7**). These data suggest R1070W may destabilize the ΔF508 structure in our models by increasing the overall score.

Since R1070W destabilized ΔF508 CFTR on a whole protein level, we sought to determine if R1070W conferred any local structural changes to the protein. We mapped the difference in residue RMSD of the lowest scoring 100 ΔF508/R1070W models vs. the lowest scoring 100 ΔF508 models onto the published closed and open conformations (**Figure 5B, 5C**). Here red represents regions unstable in ΔF508/R1070W but stable ΔF508, and blue represents regions unstable in ΔF508 but stable ΔF508/R1070W. By this metric, R1070W stabilized ΔF508 more effectively in the open state compared to the closed state (**Figure 5C**). R1070W reduced flexibility in the NBD1 α-helical subdomain of both conformations (**Supplemental Figure S8A, S8B**). These data indicate our multi-domain ΔF508/R1070W models reduced ΔF508 thermodynamic fluctuations, particularly in the open state, consistent with R1070W stabilizing effects on ΔF508 CFTR (15,16).

R1070W stabilizes the NBD1/TMD interface *in vitro* (15,16). To study this effect in our models, we calculated the residue interactions energies for each of the best scoring 100 structures and examined the energetic changes to the domain-domain interfaces. Consistent with experiment, R1070W reduces the NBD1/TMD2 interface energy compared ΔF508 alone in the closed and open conformations (**Figure 5D**). Furthermore, R1070W reduced the TMD1/TMD2 interface energy compared to ΔF508 in the open conformation (**Figure 5E**), showing a reduction towards WT levels of interface energy (**Supplemental Figure S6**). R1070W also reduced the energy of the TMD1/NBD2 interface (**Supplemental Figure S9**). These data suggest R1070W, despite having higher total potential energy in terms of Rosetta scores, primary conferred local stability in the TMD interfaces.

Our multi-domain models allowed us to simulate SSS mutations that work at NBD1/TMD interface, known to be important for ΔF508 CFTR. We simulated ΔF508/R1070W, but R1070W destabilized the structure in terms of total energy. However, R1070W stabilized the local interactions by lowering the residue RMSD of in the open state and lowering the residue interaction energy of the NBD1/TMD2 interface.

### Modeling multi-domain ΔF508 CFTR bound to VX-809

Current CF drug treatment uses small molecules called correctors to stabilize F508del CFTR including the FDA approved compound VX-809 (**Figure 6A**). Recently, two studies converged on a putative binding site for VX-809 to the WT CFTR protein (11,12). We sought to model VX-809 in our ΔF508 comparative models to determine the energetic changes VX-809 confers to ΔF508 CFTR.

**Figure 6.**
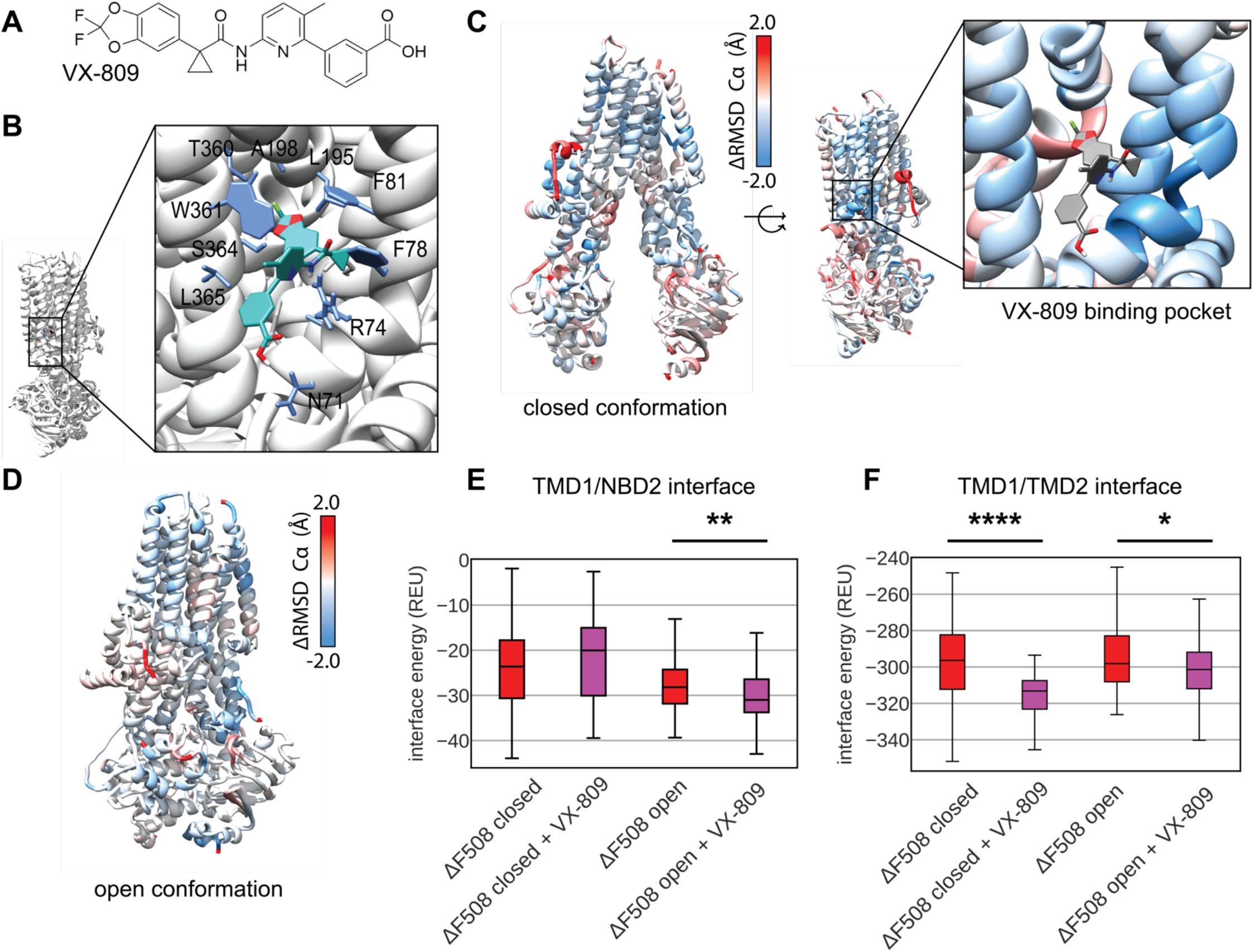
Comparative modeling of VX-809 bound to TMD1 ΔF508 reveals local stability changes including the TMD1 domain-domain interfaces. **A**. VX-809 docked to 6MSM CFTR structure in a putative binding site recently published by two parallel studies.(11,12) Interactions with important residues are shown in blue, VX-809 is shown in green with colored hetero-atoms. **B**. Average residue Cα RMSD of the lowest scoring 100 closed state ΔF508+VX-809 models subtracted from the Cα RMSD of the lowest scoring 100 closed state ΔF508 models mapped on 5UAK. Here red represents region where the RMSD was higher in ΔF508 +VX-809 than ΔF508 alone, and blue represents regions where the RMSD was lower hence the structure was stabilized by VX-809. The inset shows the RMSD of the region surrounding VX-809 demonstrates reduced RMSD. **C**. Average residue Cα RMSD of the lowest scoring 100 closed state ΔF508+VX-809 models subtracted from the Cα RMSD of the lowest scoring 100 closed state ΔF508 models mapped on 6MSM. **D**. Quantification of the residue-residue interactions at the TMD1/NBD2 interface across the lowest scoring 100 models. **E**. Quantification of the residue-residue interactions at the TMD1/TMD2 interface across the lowest scoring 100 models.

First, we docked VX-809 in Autodock to the open conformation to get an initial binding pose of non-hydrogen atoms and determine the central coordinates of the molecule. Next, we used these central coordinates to dock a full atom model into binding pocket of open conformation in Rosetta and observed the lowest scoring 10 models in terms of interface energy score. We chose a binding pose that closely resembled published cryo-EM binding site which includes interactions with W361, T360, A198, L195, F81, F78, R74, and N71 (**Figure 6B**) (11). Next, the full atom docked coordinates were copied into each ΔF508 template in the closed and open state for RosettaCM.

We generated 2000 structures of ΔF508 CFTR bound to VX-809 and analyzed the best scoring 100 models in terms of Rosetta Score. VX-809 increased the overall energy in the closed state (**Supplemental Figure S10A**), however, VX-809 reduced the overall energy in the open state (**Supplemental Figure S10B**). To look at local changes in fluctuations we mapped the difference in Ca RMSD between VX-809 bound and unbound ΔF508 CFTR onto the closed state model (**Figure 6C, Supplemental Figure S11A**). Here, blue represents areas where VX-809 reduced flexibility compared to ΔF508 alone and red represents areas where it increased flexibility. Notably, VX-809 reduce the RMSD in the binding pocket in the closed state (**Figure 6C**). We also mapped the RMSD difference onto the open state revealing VX-809 reduced flexibility in NBD1 and ICL2 (**Figure 6D, Supplemental Figure S11B**).

VX-809 allosterically stabilizes the NBD1/TMD interface *in vitro* (13,35). We calculated the residue interaction energy between the domain/domain interfaces in the presence of VX-809. We found that VX-809 increased the energy of the NBD1/NBD2 interface in our model (**Supplemental Figure S12**), however VX-809 reduced the TMD1/NBD2 interface energy in the open state (**Figure 6E**). Furthermore, VX-809 reduced the TMD1/TMD2 interface energy in both conformations (**Figure 6F**).

Thus, including VX-809 in our ΔF508 CFTR comparative models showed a reduction in overall energy in the open state. VX-809 also reduced local RMSD in the closed state around the binding site and reduced the energy in the TMD1/TMD2 interface.

## CONCLUSION

In conclusion, we used Rosetta to develop multi-domain models of ΔF508 CFTR, the primary drug target for CF. There remains a need for methods that can efficiently model large proteins, particularly proteins such as CFTR which are important drug targets. We combined cryo-EM refinement with RosettaCM to model ΔF508 and compare it to WT modeling as a control. Our models captured the thermodynamic instability of ΔF508, particularly interactions at the NBD1/TMD2 interface. These models provide a basis for computer-based drug design of CFTR correctors to target and stabilize ΔF508 CFTR.

Previous studies have used computer models to understand ΔF508 effects on the structure of NBD1. However, few studies have attempted to understand ΔF508 effects on multi-domain CFTR, despite the recent publications of these cryo-EM structures. ΔF508 folding defects stem from structural defects in both NBD1 and the NBD1/TMD interface, and ΔF508 correction requires fixing both defects (15,16). Hence, it is imperative to develop multi-domain models of CFTR to gain insight into the atomic level interactions underlying these structural defects. Here we used comparative modeling in Rosetta (RosettaCM) as it offers the computational speed required for virtual drug screening

We used recently published cryo-EM models of full length CFTR – with an undetermined R domain – to refine an ensemble of WT models in the closed and open conformations. The lowest scoring models from refinement were used as templates for RosettaCM by threading the ΔF508 sequence onto the structure. We tested the number of templates required to capture ΔF508 thermodynamic instability by simulating 1000 models of WT and ΔF508 NBD1 with and without SSS mutations with known ΔΔG and ΔT_M_ values in the literature. Using five cryo-EM refinement models correlated the best with experimental data. We then applied this sampling method to multi-domain ΔF508 CFTR.

Here, we developed a RosettaCM approach for modeling ΔF508 CFTR compatible with RosettaLigand and the rest of the Rosetta Software Suite we will leverage for future rational CFTR drug design. Our multi-domain models are still missing many loop regions that remain undetermined in the cryo-EM density. For example, the regulatory insertion (RI) region changes the thermodynamic stability of CFTR (36) and adopts distinct conformations, one of which has been postulated to lead to ΔF508 unfolding (37). Our models are also missing the R domain (a large unstructured 200 residues between NBD1 and TMD2), the glycosylation site, and the loop connecting TMD2 to NBD2. Modeling loop regions either with loop modeling in Rosetta or using the cryo-EM density and Rosetta Enumerated Sampling will further improve the biological relevance of our approach.

Our multi-domain ΔF508 models presents advantages and limitations towards the goal of providing a basis for computer-based drug design. The closed and open state ΔF508 models successfully captured the thermodynamic instability of ΔF508 CFTR evident by the overall higher Rosetta scores of these models compared to WT. However, R1070W destabilizes the protein structure in our models, but stabilized the NBD1/TMD2 interface suggesting that our models captured local energetic changes but failed to capture global changes.

Corrector compounds stabilize both ΔF508 NBD1 and the NBD1/TMD interface. Thus, modeling multi-domain ΔF508 CFTR represents a key step towards structure-based drug design for CF. We found, among the currently available multi-domain CFTR cryo-EM structures, modeling the open conformation to be consistent with known experimental ΔF508 instability. VX-809 stabilized ΔF508 CFTR in the open state when included in our model. Thus, the open state and may offer more biologically relevant sampling with this technique than the closed state. The ΔF508 CFTR and VX-809 modeling approach developed here will aid future rational, structure-based drug development efforts for CF.

## METHODS

### Protein Structural Data Preparation

The dephosphorylated (closed) human CFTR cryo-EM structure was downloaded from the PDB (5UAK (2), resolution 3.9 Å, determined residues 5-402, 439-645, 845-883, 909-1172, 1207-1436). The residues from a poorly determined helix between the TMDs were removed from the 5UAK PDB file manually. We also downloaded the phosphorylated (open) human CFTR cryo-EM structure from the PDB (6MSM (20), resolution 3.2 Å, determined residues 1-409, 435-637, 845-889, 900-1173, 1202-1451) and removed lipids and an unresolved helix near the lasso motif manually, ATP was kept in both binding sites. 6MSM contains the stabilizing mutation E1371Q and we used the MutateResidue mover in Rosetta to revert this back to E in our model. Finally, we also downloaded the raw cryo-EM density maps for 5UAK and 6MSM from the PDB.

For our testing set, we prepared a NBD1 structure from 5UAK by truncating the published model at residue Y385 through the determined portion of NBD1 to residue M645 (note this excludes the RI region from 403-438). We modeled ATP into the degenerate site by aligning 5UAK and 6MSM and copying the MG and ATP coordinates from 6MSM into the NBD1 structure. This resulted in an NBD1 structure including ATP bound at the degenerate site for our testing set (**Figure 3B**).

### Cryo-EM Refinement

We refined the published coordinates into the raw cryo-EM density maps using a previously established method in Rosetta (23) (see Protocol Capture Step 1). This approach requires the published structure coordinates and the published cryo-EM density map (both available on the PDB) as well as a set of refinement parameters (**Supplemental Figure S1**). We optimized refinement parameters including the weight put on the cryo-EM density, the length of fragment insertion and the distance of fragment insertion to increase ensemble diversity. We evaluated ensemble diversity by calculating the structural alpha carbon (Cα) root mean squared deviations (RMSD) from the published model for each refinement ensemble and assumed a greater Cα RMSD distribution indicated a more diverse ensemble. We optimized the sampling weight put on the cryo-EM data (denswt), the root mean squared distance (RMS) for peptide fragment insertion, and the length of the peptide fragments (**Supplemental Figure S1**).

### Optimization of Cryo-EM Refinement Parameters

We optimized the user specified parameters required for Rosetta cryo-EM refinement (23). First, to avoid overfitting, we optimized the weight put on the experimental density data (denswt) in the refinement score function. We generated 100 structures at density weight values between 20 and 50 at 5-point intervals. Next, we plotted these density weight values versus the difference between the Fourier Shell Correlation (FSC) and 4% of the per-residue energy for the ensemble (FSC – 0.04*per-residue energy). The maximum difference indicates the optimal density weight. We chose a density weight value of 30 as this maximizes the (FSC – 0.04*per-residue energy) value for most structures (**Supplemental Figure S1A, S1B**). Second, to maximize structural diversity, we optimized the length of peptide fragment insertion. The refinement protocol builds possible models by breaking sequences of determined residues into peptide fragments of an odd number length (e.g. 5, 7, 9, 11, or 13) (23). Increasing the insertion length increases model diversity (23). We increased the fragment insertion length from seven to thirteen, generated 100 models for each, and plotted the Cα RMSD of each model in the refinement from the published model versus the model score for all four CFTR structures. Indeed, increasing the fragment insertion length from seven to thirteen generated overall lower scoring models with a greater Cα RMSD distribution for both the closed (PDB ID 5UAK) and open (PDB ID 6MSM) states (**Supplemental Figure S1C, S1D**). Thus, we chose 13 for our fragment length. Third, we optimized the root mean squared (rms) distance between the inserted peptide fragments by varying this value from 1.5 Å to 2.5 Å in intervals of 0.25 and generating 100 models per interval. We plotted the Cα RMSD of the most poorly determined domain - NBD1 - vs. the model score. We plotted NBD1 as this domain will likely have the greatest distribution in structural diversity from refinement. We chose a rms value of 1.75, as this value increases the NBD1 Cα RMSD (**Supplemental Figure S1E, S1F**). Increasing the rms value beyond 1.75 offered no improvement in NBD1 Cα RMSD (data not shown for clarity).

### *In Silico* Mutagenesis

We made point mutations in CFTR structures using the MutateResidue mover in Rosetta. For the phosphorylated model, 6MSM, we mutated E1371Q back to the naturally occurring glutamine residue. The low structural resolution makes side chains difficult to distinguish in regions of NBD1, near F508. Hence to generate deletion mutations, we removed F508 from the open and closed state CFTR fasta files respectively and threaded the sequence onto the open and closed state models. For our testing NBD1 structure, we again deleted F508 from the NBD1 fasta sequence (residue 385-402 and 439-645) and threaded the new sequence onto the NBD1 structure. We mutated all second site suppressor mutations (F494N, F494N/Q637R, V510D, I539T, and G550E/R553Q/R555K) in NBD1constructs prepared for our testing set using the MutateResidue mover in Rosetta.

### Rosetta Comparative Modeling

To model CFTR variants, we used RosettCM, a homology modeling approach(24). We perform CM with static templates derived from the cryo-EM density, not the cryo-EM density itself. Published CFTR structures contain undetermined loops and intrinsically disordered regions including the RI region, the RD, the glycosylation site, and the loop linking TMD2 to NBD2. We generated fasta files containing only the determined residues in 5UAK and 6MSM (**Supplemental Table S2**). To generate ΔF508 templates, we manually removed F508 from the fasta files and threaded the ΔF sequence onto the five WT models (see *in silico* mutagenesis section). As a control we used the WT sequence and the original WT templates and performed the same modeling protocol. Additionally, we mutated R1070W into the ΔF508 templates and substituted the W manually to the fasta sequence to model ΔF508/R1070W CFTR.

We performed multiple template hybridization with the Hybridize mover in Rosetta guided by the RosettaMembrane energy function (38,39). We imposed membrane specific Rosetta energy terms within the theoretical membrane bilayer by predicting the transmembrane helix regions with OCTOPUS (40). We set all template weights to 1.0. For fragment insertion, we used three and nine peptide long fragments with short and long fragment insertion weights set to 1.0. We optimized side chain positions by simulated annealing, also known as rotamer packing in Rosetta. We refined final models using one round of FastRelax mover (e.g. repeat =1) in Rosetta which performs steepest gradient decent minimization in Cartesian coordinate space without constraints.

### Calculation of Protein Stability Metrics

We evaluated protein thermodynamic stability metrics for WT, ΔF508, and ΔF508/R1070W CFTR. We calculated the alpha carbon (Cα) root mean squared deviation (RMSD) for whole structures as well as on a per-residue basis with respect to a reference model (either the published model or a low scoring model in the ensemble). For our per-residue Cα RMSD calculations we first aligned individual domains to account for any shifts in the domains relative to each other as we were interested only in local fluctuations. We assumed local Cα RMSD as a surrogate for protein flexibility. Furthermore, we calculated the residue interaction potential energy, which provides the potential energy in REU between every pair of contacting side chain in the structure. Finally, we calculated Rosetta scores for our NBD1 testing with second site suppressor mutations with ref2015 and calculated Rosetta scores using the membrane scoring function (38,39).

### Docking and Parameterizing VX-809

An automated docking tool, Autodock Vina (41) version 1.1.2, Scripps Research Institute (La Jolla, CA), was employed to dock parameterized VX-809 on the CFTR open cryo-EM structure (PDB-ID: 6MSM). Top 20 binding modes in the energy range of 100 kcal/mol were computed. Grid parameters: center (x, y, z) = (165.453, 168.303, 149.074), size (x, y, z) = (20, 20, 20). To get the full atom parameters for VX-809, we created low energy 3-dimensional ligand conformations in Corina given 2D representation exported from Chem draw. We then checked the BCL-based basic chemistry for appropriate bond lengths, atom types, etc. Next, we generated ligand conformers using BCL ConformerGenerator (42) for 8000 iterations and clustered based on distance between individual conformers. We then made Rosetta-readable parameters file for ligand docking and comparative modeling. This takes the conformer SDF and assigns partial charges and points to the conformer file. This also outputs centroid and torsional parameter files, which are used in the comparative modeling with CFTR. We then performed full atom docking in Rosetta using RosettaLigand (43).

## ACKNOWLEDGEMENTS

We thank members of the Meiler lab for their critical reading and feedback of this manuscript.

## AUTHOR CONTRIBUTIONS

EFM, HW, LP, and JM conceived the study and planned the simulations. EFM prepared structural models, set up simulations, and performed simulation. EFM and CTS analyzed the data. EFM prepared the figures and wrote the manuscript. SS and MK generated VX-809 models, docked models, and performed parameterization. LP, JM, HW, and MK critically reviewed the manuscript. JM and EFM edited the manuscript.

## DATA AVAILABILITY

A protocol capture on how to generate CFTR refinement models as well as CFTR comparative models is included in the Supplemental Information.

